# A canine brain bank for comparative neuroscience and brain aging research

**DOI:** 10.64898/2026.07.11.737944

**Authors:** Sarah Darcy, Autumn Beck, Macy Garrood, Andria Slaughter, Alexander Parra, Laura Paredes, Kurt Farrell, John F. Crary, Andrew T. McKenzie

**Affiliations:** Apex Neuroscience, Salem, OR, 97317, USA; Friedman Brain Institute, Departments of Pathology, Neuroscience, and Artificial Intelligence & Human Health, Icahn School of Medicine at Mount Sinai, New York, New York, USA; Neuropathology Brain Bank & Research Core and Ronald M. Loeb Center for Alzheimer’s Disease, Icahn School of Medicine at Mount Sinai, New York, New York, USA

**Keywords:** Brain banking, Perfusion fixation, Canine cognitive dysfunction, Lipofuscin, Postmortem interval

## Abstract

Companion animal brain banking has been recognized as a valuable approach for translational aging and dementia research. However, realizing the full value of canine brain banks depends on optimizing the methods that are used to collect and preserve the tissue. Whole brain perfusion fixation is one promising approach, but it is not yet well described in dogs. Here we describe the development of methods for a canine brain bank (currently n = 55), including whole brain perfusion fixation via aortic cannulation and brain extraction. We assessed perfusion quality using gross examination, post-perfusion CT, and histological clearance of blood vessels. We found that body weight and average flow rate per body weight were each significantly correlated with perfusion quality in our cohort. To illustrate the kind of analysis the bank could facilitate, we next performed a preliminary study of brain aging, one of our primary planned research applications. Using a pixel classifier applied to whole slide images, we quantified lipofuscin burden, and in this preliminary cohort found that it increased strongly with age in both the thalamus and hippocampus. In the hippocampus, lipofuscin burden was also elevated in dogs with owner-reported cognitive dysfunction, although the current cohort is too small to determine to what extent this association is independent of age. Preliminary electron microscopy studies also confirmed that perfusion fixed tissue from the bank is amenable to ultrastructural analysis. This work describes one approach for canine brain perfusion fixation and introduces a brain tissue resource that may help support future neuroscience research.

## Introduction

Brain banking is an essential component of the neuroscience research community to promote the study of the brain in both health and disease. The majority of brain banks focus on human brains. However, in recent years, companion animal biobanking has gained increasing attention, and multiple canine brain banks have now been established in Europe and North America (Sándor et al., 2021; Cardy et al., 2022; McEnhill et al., 2024; McGrath et al., 2025). There are several advantages to banking companion animal brains, including their value as naturally occurring models of neurobiological diseases, and the potential for comparative neurobiology across species to illuminate general principles of brain function and dysfunction. There is growing recognition of the potential value of dogs in particular as a translational model for Alzheimer’s disease and related dementias (Head, 2013). This has motivated the development of at least one large-scale longitudinal canine brain banking study (McGrath et al., 2025). Beyond their relevance to disease modeling, humans are naturally curious about canine brains, given the remarkable ability of dogs to cooperate with and understand humans, which is of course dependent upon their brain structure and its modification through domestication (Barton et al., 2025). In the long term, canine brain banks can contribute to both of these goals, helping to facilitate studies of brain aging and neurodegeneration and also shedding light on the neural basis of the unique cognitive and social capacities of dogs.

For preserving the structure of an entire mammalian brain, perfusion fixation is considered the gold standard method (Bodian, 1936; McFadden et al., 2019). While there are many perfusion fixation protocols available for laboratory animals (Gage et al., 2012; Wahyudi et al., 2025), to the best of our knowledge, there is not an extensive literature on the perfusion of canine brains. Canine cerebrovascular anatomy presents unique considerations. The brain is supplied in part by a number of anastomoses between the intracranial circulation and the external carotid artery (Jewell, 1952; Lee et al., 1986). The most important of these is called the anastomotic artery, which forms from a branch of the internal maxillary artery – and in some cases also the middle meningeal artery – and connects to the internal carotid artery (Jewell, 1952). In addition, canine cerebrovasculature is posterior circulation dominant. As a result, contrast injected into the basilar artery leads to visualization of bilateral territory throughout the brain, whereas contrast injected into the internal carotid artery leads only to the visualization of the territory of the middle cerebral artery and (less reliably) the anterior cerebral artery (Camstra et al., 2020). Finally, many of the arteries supplying the canine brain are highly tortuous, which may raise the risk of resistance to perfusate flow, based on reports of perfusion fixation in other animals (Eden and Correia, 1981).

As they age, some dogs develop a syndrome known as canine cognitive dysfunction, which shares clinical and pathological features with human Alzheimer’s disease, including amyloid beta deposition, neuronal loss, and behavioral deficits (Cummings et al., 1996; Head, 2013). One of the most consistent cellular correlates of brain aging, in both dogs and humans, is the accumulation of lipofuscin. Lipofuscin is an autofluorescent pigment composed of oxidized lipid and protein residues that accumulates in the lysosomes of postmitotic cells and causes a variety of deficits in cellular functioning (Brunk and Terman, 2002). Lipofuscin has been found to accumulate progressively in canine neurons during aging, with differences between regions in the timing of onset and rate of accumulation (Whiteford and Getty, 1966; Few and Getty, 1967; Nanda and Getty, 1973; Nesic et al., 2021). Extensive lipofuscin accumulation has also been associated with canine cognitive dysfunction in one case report (Park et al., 2016). However, to the best of our knowledge, the extent to which lipofuscin accumulation in the canine brain is quantitatively associated with canine cognitive dysfunction in larger cohorts, as opposed to simply correlating with chronological age, is an open question.

In this manuscript, we describe our canine brain bank, methods for perfusion fixation via aortic cannulation, and our preliminary assessment of perfusion quality using gross examination, CT neuroimaging, light and electron microscopy. Among the histological findings, we focus particularly on lipofuscin, which we quantify using a pixel classifier applied to whole slide images and examine in relation to both age and canine cognitive dysfunction. We also describe our future plans and directions for the canine brain bank. Our primary aim here is to describe and document the methods we developed, and the histological and ultrastructural analyses we present are intended as examples of the types of research applications that the cohort could support in the future.

## Methods

### Canine donors and tissue collection

We built upon previous efforts in canine brain banking to create our own canine brain bank (Sándor et al., 2021; McGrath et al., 2025). We partnered with local veterinary professionals to provide euthanasia services in our facility for canine companion animals whose owners elected this humane end-of-life care option. This was done in order to minimize the postmortem interval (PMI) and to provide a benefit to companion animal owners by covering the cost of euthanasia services. Because all dogs were companion animals undergoing owner-elected euthanasia for end-of-life care, independent of this study, and brain tissue was obtained postmortem with owner consent, no animals were euthanized or otherwise used for research purposes. The work therefore did not require Institutional Animal Care and Use Committee approval.

For a basic assessment of whether cognitive dysfunction was present, owners were asked “Has your pet shown signs of confusion in the past six months, such as getting lost or having difficulty recognizing familiar people?” with possible answers of “No,” “Mild,” “Moderate,” or “Severe.” For the analysis comparing lipofuscin burden by cognitive status, because the small sample size precluded a graded ordinal analysis, responses were dichotomized into absent (No) versus present (Mild, Moderate, or Severe).

For each case, the PMI was calculated as the time between cardiac arrest and the onset of perfusion. In some cases, the PMI was estimated based on known sequences and duration of procedural steps, rather than precise timestamps, due to inconsistent data collection about the timing of euthanasia and the procedure.

### Perfusion methods

Our perfusion methods built upon previous techniques that have been used in human whole body donors (Garrood et al., 2025b). They were improved upon in an iterative manner. In early experiments, we performed cannulation of the carotid arteries. This was found to be ineffective for perfusion of the whole brain in most cases, which we attributed to differences in cerebrovascular anatomy between humans and dogs (Jewell, 1952). Another method we attempted was transcardial perfusion via the left ventricle, but this method was also discontinued due to inconsistent perfusion quality and difficulty in securing the cannula. The method that was found to produce the highest quality of perfusion with the most consistency was perfusion via cannulation of the aorta, for which we describe our current working protocol.

Following humane euthanasia by a licensed veterinarian, a thoracotomy is performed and a sternal retractor is used to open the chest cavity. Throughout the procedure, an aspirator is used to manage blood loss and keep the cavity clear of excess fluid. Once the chest cavity is opened, the pericardium is incised and reflected from the heart and surrounding structures using forceps and dissecting scissors, which is generally necessary in order to visualize the aorta. Variations in mediastinal anatomy (e.g., differences in adipose tissue, adhesions, or occasional masses) between dogs of different breeds or health histories are a common occurrence. Effectively clearing visceral tissue is essential for accurate identification of the aorta, and this needs to also be done rapidly to minimize the PMI, while also accurately to avoid incising the aorta or any other vessels.

After clear visualization of the proximal aorta, we place a Satinsky DeBakey clamp across the proximal aorta, partially occluding the vessel and retracting the heart caudally to expose a larger segment for cannulation. A scalpel with a #11 blade is used to make a puncture in the anterior aortic wall, taking care not to puncture the opposite wall. The cannula is then inserted into the incision and held manually in place throughout the perfusion. The cannula has a raised ring above the tip that rests against the outer aortic wall, limiting insertion depth and helping to maintain a consistent cannula position. Manual stabilization is used to ensure that the cannula tip stays angled toward the aortic arch and prevent decannulation as flow rate and/or perfusion pressure increases during the course of the perfusion. Occasionally, leakage occurs around the cannulation site, in which case hemostats can be used to clamp the tissue surrounding the cannula to establish or re-establish a seal. This approach is feasible given our relatively short perfusion duration and is analogous to the initial phase of cannula stabilization described in one protocol for normothermic regional perfusion in donation after circulatory death organ procurement, in which the cannula is held manually until stable flow is established (Trahanas et al., 2022).

Following the placement of the curved tip aortic cannula (22 Fr), the circuit is de-aired and the perfusion phase of the procedure is initiated. Perfusion is driven by a peristaltic pump system. Pressure is monitored using an inline pressure sensor connected to the perfusion circuit. The right atrial appendage is cut for outflow as blood and perfusate fluid return to the heart via the superior vena cava. An aspirator is used to suction the fluid in the thoracic cavity and keep the surgical field clear. The peristaltic pump speed was gradually increased throughout the early stages of the perfusion procedure until a desired pressure was reached, of approximately 100-150 mmHg. The highest pressure throughout the procedure was also recorded and occasionally was higher than this. The maximum flow rate was determined case by case, depending on the pressure. Fluid return during perfusion was monitored where the right atrium was cut. In most cases, a gradual transition of the color of return fluid from dark red to clear or mostly clear was observed. In larger dogs, we found that it was necessary to perfuse a larger volume of fluid in order to drain the blood.

Batches of perfusate were prepared in polycarbonate carboys. In most cases, the base solution was 10% or 20% neutral buffered formalin (NBF). Depending on the case, we added one or more of the following: glutaraldehyde at 1% (v/v), to attempt to improve ultrastructural preservation; iohexol at 3 mg/mL, for contrast-enhanced CT imaging; green dye, to better visualize color changes in brain tissue on gross examination; and mannitol at 10% or 20% (w/v) and/or polyethylene glycol 35 kDa (PEG35) at 5% or 10% (w/v), to increase osmotic or oncotic pressure in an attempt to improve ischemic perfusion quality. Carboys were stored at 4°C or at room temperature until use.

When the CT scanner was available, CT scanning was performed after the perfusion, with the brain still inside the skull. The CT scanner we used was the OmniTom® Elite (Neurologica, Danvers, MA), a 16-slice scanner. Images were viewed with the Osimis Web Viewer.

**Table 1.**
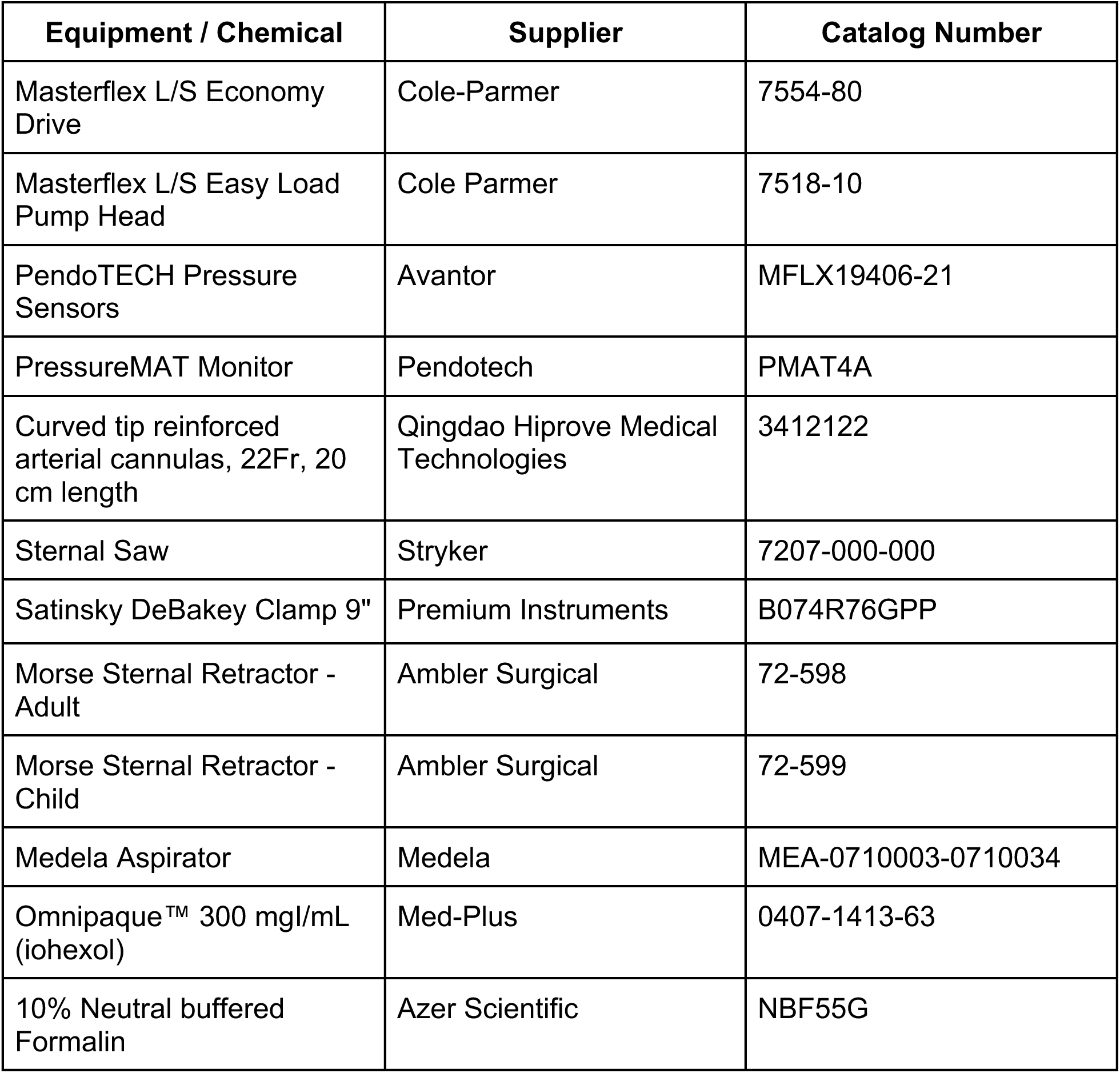
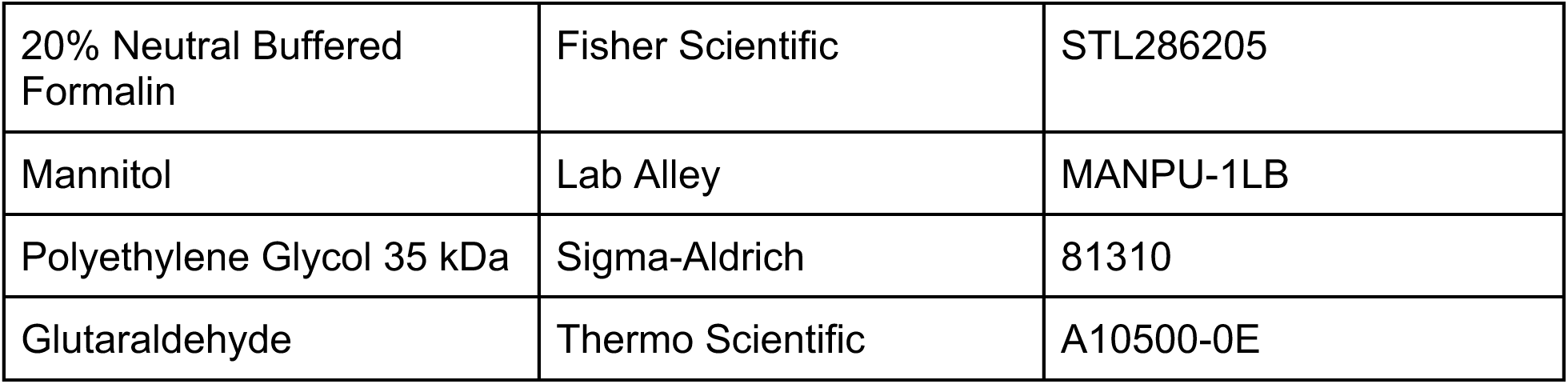
Supplies and reagents used for perfusion fixation.

| Equipment / Chemical | Supplier | Catalog Number |
| --- | --- | --- |
| Masterflex L/S Economy Drive | Cole-Parmer | 7554-80 |
| Masterflex L/S Easy Load Pump Head | Cole Parmer | 7518-10 |
| PendoTECH Pressure Sensors | Avantor | MFLX19406-21 |
| PressureMAT Monitor | Pendotech | PMAT4A |
| Curved tip reinforced arterial cannulas, 22Fr, 20 cm length | Qingdao Hiprove Medical Technologies | 3412122 |
| Sternal Saw | Stryker | 7207-000-000 |
| Satinsky DeBakey Clamp 9" | Premium Instruments | B074R76GPP |
| Morse Sternal Retractor - Adult | Ambler Surgical | 72-598 |
| Morse Sternal Retractor - Child | Ambler Surgical | 72-599 |
| Medela Aspirator | Medela | MEA-0710003-0710034 |
| Omnipaque™ 300 mgI/mL (iohexol) | Med-Plus | 0407-1413-63 |
| 10% Neutral buffered Formalin | Azer Scientific | NBF55G |
| 20% Neutral Buffered Formalin | Fisher Scientific | STL286205 |
| Mannitol | Lab Alley | MANPU-1LB |
| Polyethylene Glycol 35 kDa | Sigma-Aldrich | 81310 |
| Glutaraldehyde | Thermo Scientific | A10500-0E |

### Brain extraction method

Brains were extracted according to the following protocol, applied with or without prior perfusion fixation. The extraction protocol was refined iteratively over the course of the study and we describe our current working approach below. It was primarily adapted from the protocol we use for human brain extraction (Wolf et al., 2026).

First, the skin and musculature overlying the calvarium were dissected and reflected to expose the underlying bone. An oscillating bone saw was then used to perform the craniotomy, applied in a light, smooth, reciprocating motion to avoid penetrating the full thickness of the skull and damaging the underlying neural tissue. The cut was initiated on the lateral aspect of the frontal bone and carried across the parietal bone at the widest point of the cranium. This incision was continued circumferentially, passing inferior to the occipital protuberance and returning along the contralateral frontal bone, with the two cut lines converging at a V-shaped angle along the frontal bone and forehead of the specimen. This cut produces a triangular window exposing the frontal sinus cavity, providing the access required to isolate and remove the olfactory bulbs.

Because skull morphology and thickness vary among dog breeds, some specimens required additional force, applied with an osteotome and mallet advanced incrementally along the entire saw cut line, to release the dorsal portion of the cranium. In certain cases, the V-shaped cut was repeated on the newly exposed internal table of the frontal bone. The craniotomy was completed by inserting a skull breaker along the saw cut line and twisting it to fully release each margin of the calvarium. In especially difficult cases, an additional sagittal cut was made through the midline and the cranium was removed in multiple pieces.

Following removal of the calvarium, the dura mater was incised and reflected away from the brain. Isolation of the olfactory bulbs was then performed by clearing the sinus tissue from the sinus passage with forceps and removing any residual fragments of the internal table of the frontal bone overlying the bulbs using side-cutting rongeurs. Once the bulbs were exposed, a micro-spatula was used to gently sweep around and detach them from the cribriform plate of the ethmoid bone. The brain, including the cerebellum, was subsequently freed by passing a malleable brain spatula smoothly around its perimeter within the cranial vault. Once sufficiently loosened, the brain was gently pulled posteriorly to expose and transect the optic nerves, then gently pushed anteriorly to expose and cut the spinal cord. The brain was then lifted out of the cranial vault, with care taken to ensure that the olfactory bulbs remained attached to the brain and were not torn away at the ethmoid bone.

In cases where the brain appeared soft following skull removal, the entire cephalon was occasionally placed in a container of 10% NBF to allow for *in situ* fixation prior to brain extraction the following day or up to several days later. This minimizes damage to the brain during the extraction process when there is inadequate perfusion fixation (or it was not performed at all). After extraction, digital photographs were taken of the brain from multiple angles for gross assessment. After the photographs were taken, the brains were completely immersed in a plastic container with 10% NBF, which was placed at 4°C for long-term preservation prior to histological assessment (McKenzie et al., 2024).

**Table 2.** Supplies used for brain extraction.

| Equipment | Supplier | Product Identifier |
| --- | --- | --- |
| Scalpel Handle No 3 | MedHelp | B08QDMFJRG |
| Feather Scalpel Blade #22 | Electron Microscopy Sciences | 72044-22 |
| Mopec 810 Autopsy Saw | Mopec | BD810 |
| Standard Small Section Pathology Saw Blade | Mopec | BD108 |
| Lexer Bone Osteotome 15 mm x 22 cm with Fiber Handle | Medikrebs | TT283 |
| Heavy Bone Mallet | Surgical Online | B07CRQTXDW |
| Brain Spatula, Malleable, 16 mm x 20 cm | Medikrebs | B07QW3MZH6 |
| Skull Breaker, 3" | Integra | Miltex 34-210 |
| Ferris Smith Tissue Forceps, 7 inch, Serrated Platform Tips with 2 x 3 Teeth | Integra Miltex | 26-958 |
| Standard Surgical Scissors, Blunt/Blunt, Curved | Fine Science Tools | 14003-13 |
| Medium curved or straight rongeurs (14-16 cm) | Fine Science Tools | 16120-14 |
| Micro-spatula | Fine Science Tools | 10091-12 |

### Light microscopy

In a subset of 21 cases, we sectioned the brains and dissected approximately 1 × 1 × 1 cm tissue samples targeting the approximate hippocampus and thalamus regions for light microscopy. As a control, samples were also taken from two brains that were immersion fixed rather than perfusion fixed. Brain tissue sampled for light microscopy was placed into cassettes for processing and embedded in paraffin. Paraffin-embedded brain sections 6 μm thick were baked, deparaffinized, and stained for hematoxylin and eosin (H&E). Digital images of the stained sections were captured at 40X as whole slide images (WSIs) using the Aperio GT450 high-resolution scanner (Leica Biosystems).

### Perfusion quality assessment

Semi-qualitative analysis was performed of images from gross examination, CT imaging, light microscopy, using methods previously described and validated as internally consistent (Garrood et al., 2025b). Gross examination and CT images were graded by the extent of perfusion in eight vascular territories, i.e. the territory supplied by the left and right anterior cerebral artery (ACA), middle cerebral artery (MCA), and posterior cerebral artery (PCA), as well as the cerebellum. For gross examination, the features used for this assessment were (a) clearance of blood from surface vessels, (b) apparent stiffness of tissue, and (c) color change of tissue, from pink in the case of no perfusion to pale or pale/colored dye-tinged in the case of well perfused tissue. For CT images, grading was based on the extent of contrast visualized in post-perfusion scans. LM images were graded by the extent of vessel clearance, which refers to the absence of intravascular material from both small and large vessels. All images were graded on a scale of 0-3, with 0 indicating <5% extent of perfusion, contrast or vessel clearance, 1 indicating 5-50%, 2 indicating 50-95%, and 3 indicating >95%. Gross examination and CT images were graded by a single trained rater using these previously validated methods. Light microscopy images were graded independently by two independent raters. For light microscopy, inter-rater reliability between two raters was assessed via quadratic-weighted Cohen’s kappa, which was found to be 0.80 (indicating substantial agreement). Discrepancies were resolved for downstream analysis by consensus review.

For each modality, a composite perfusion quality score was calculated as the mean of the available regional grades (i.e. the eight vascular territories for gross and CT examination, and the hippocampus and thalamus for light microscopy). These composite quality scores were calculated only for those cases with at least four of the eight regional grades available for gross and CT, and with both regional scores available for histology. The average flow rate per body weight was calculated as the total perfusate volume in mL divided by the duration of perfusion in minutes divided by the body weight in pounds. Associations among donor characteristics, perfusion parameters, and perfusion quality metrics were assessed using Spearman rank correlations. Given the exploratory nature of these analyses, p-values are reported as nominal values without correction for multiple comparisons.

### Lipofuscin

Lipofuscin burden was quantified in each WSI using a pixel classifier trained in QuPath (v. 6.0). The classifier was a random trees pixel classifier implemented via OpenCV, using the red, green, and blue channels smoothed with a Gaussian filter (sigma = 1.0) as input features, applied at a resolution of 2.1 µm per pixel in 512 x 512 pixel tiles. Each pixel was classified as either lipofuscin-positive or background. The largest number of false positives was pigment found near blood vessels, which is presumed to be hemosiderin from blood cell breakdown products. The training annotations were generated primarily on the thalamus WSI from donor 229, which was selected on an initial qualitative review to be a representative WSI with a substantial lipofuscin burden. Tissue edges were excluded from analysis, as they also had high false positives, which may be due to artifacts occurring during processing. The trained classifier was then applied to all available WSIs within a standardized bounding box (approximately 1,000 μm x 1,000 μm), centered to avoid tissue edges and ensure a consistent size of analysis across WSIs. Lipofuscin burden was quantified as the percentage of pixels within the standardized bounding box classified as lipofuscin-positive, which we refer to as lipofuscin density. We also recorded the number of discrete lipofuscin-positive objects within the box.

Associations between age and lipofuscin measures (density and object count, in the thalamus and hippocampus) were assessed using Spearman rank correlations. Differences in lipofuscin measures between dogs with and without owner-reported cognitive dysfunction were assessed using the Mann-Whitney U test. Given the exploratory nature of these analyses, p-values are reported as nominal values without correction for multiple comparisons.

### Electron microscopy

Sample preparation and imaging was performed as previously described (Garrood et al., 2025a). Tissue for EM was post-fixed in a solution of 2 % paraformaldehyde and 2.5 % glutaraldehyde in 0.1M sodium cacodylate buffer. The tissue then underwent a multi-step enhanced contrast protocol at room temperature including sequential treatments with tannic acid, reduced osmium, thiocarbohydrazide, osmium, and uranyl acetate. This was followed by lead aspartate staining at 60 °C. The brain sample was then dehydrated through a graded ethanol series, infiltrated with Embed 812 epoxy resin (EMS), and polymerized for 72 h at 60 °C. Semithin sections (0.5 μm) were cut using a Leica UC7 ultramicrotome (Leica, Buffalo Grove, IL) and counterstained with 1% toluidine blue to identify the regions of interest within layers. Images were taken on a HT7500 transmission electron microscope (Hitachi High-Technologies, Tokyo, Japan) using an AMT NanoSprint12 12-megapixel CMOS TEM Camera System, software version 7.0.1.485 (Advanced Microscopy Techniques, Danvers, MA). Images were adjusted for contrast on the AMT software.

## Results

### Description of our cohort

Our canine brain bank currently has n = 55 donors with a wide variety of ages, sizes, and breeds (**Table 3**). Representative gross and CT images from three donors illustrate the anatomical diversity (**Figure 1**). Donor ages were owner-reported and ranged from 1 to 26 years (median 13 years), with the oldest value an outlier that may be inaccurate (the next highest was 18). We used all owner-reported ages rather than excluding certain values based on plausibility judgments. Body weights ranged from 4 to 120 lbs, capturing most of the size spectrum of the domestic dog. Using a previously described size classification based on body weight, our cohort consists of 20.00% Toy (under 11 lb), 21.82% Small (11 to under 22 lb), 14.55% Medium (22 to under 55 lb), 30.91% Large (55 to under 88 lb), and 12.73% Giant (88 lb and above) breeds (Salt et al., 2017).

**Figure 1.**
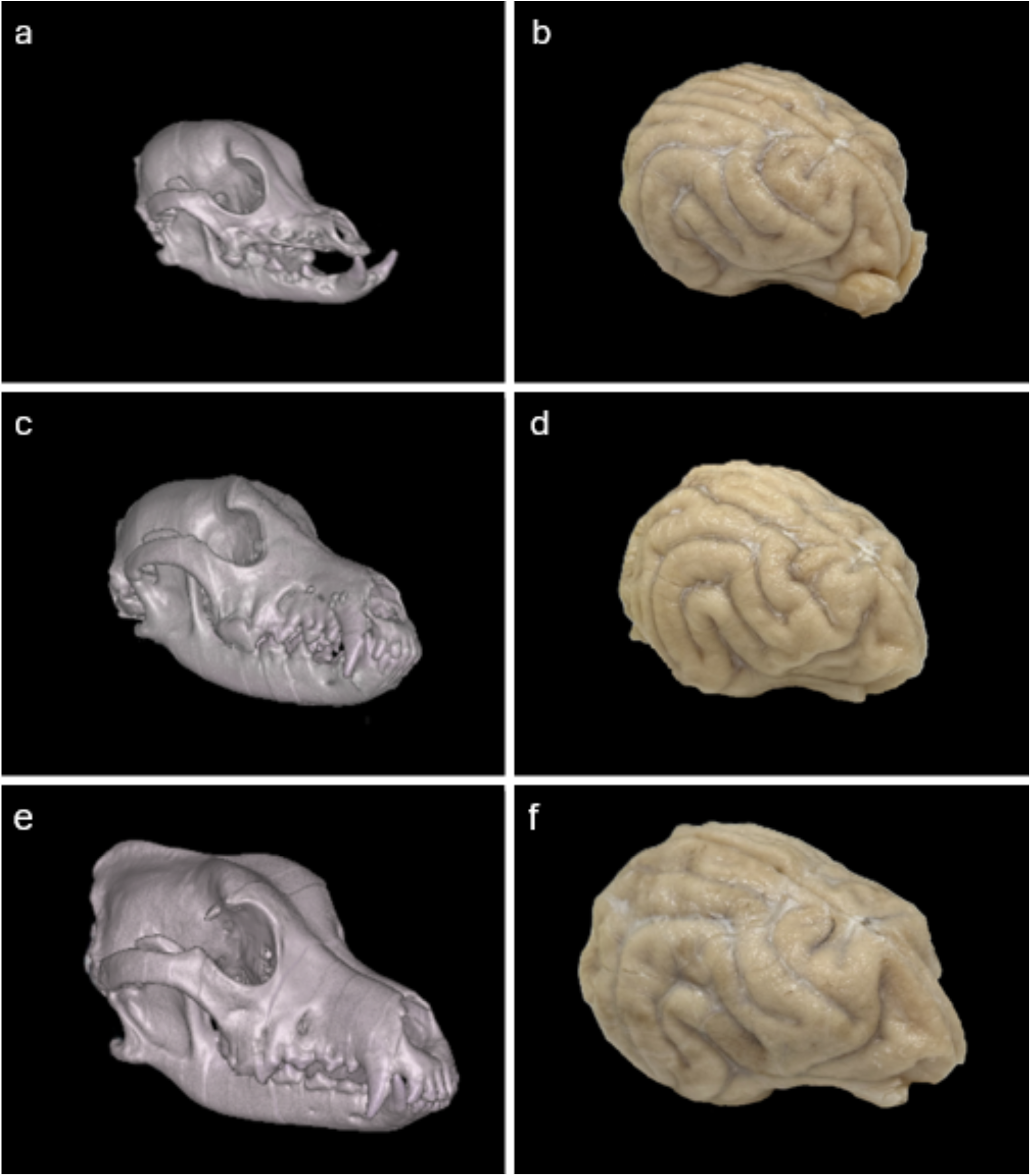
Representative gross and CT images illustrating the anatomical diversity of the canine brain bank. CT images of the head (**a**, **c**, **e**) and corresponding gross brain photographs taken after perfusion fixation and subsequent immersion fixation (**b**, **d**, **f**) are shown for three donors: donor 329 (Chihuahua–Terrier, 14 lb; **a**, **b**), donor 303 (Australian Labradoodle, 25 lb; **c**, **d**), and donor 310 (Labrador–Heeler, 60 lb; **e**, **f**).

**Table 3.**
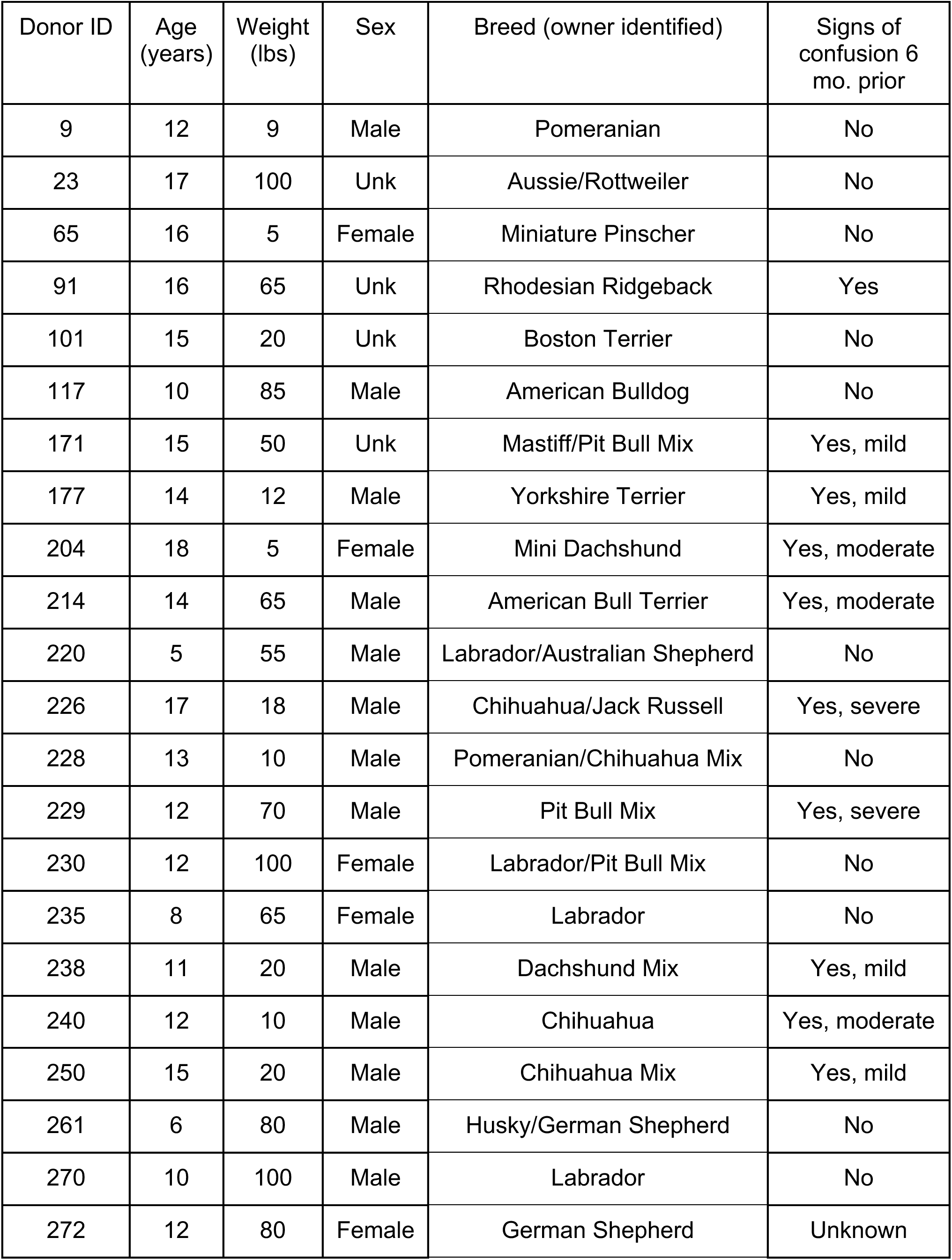
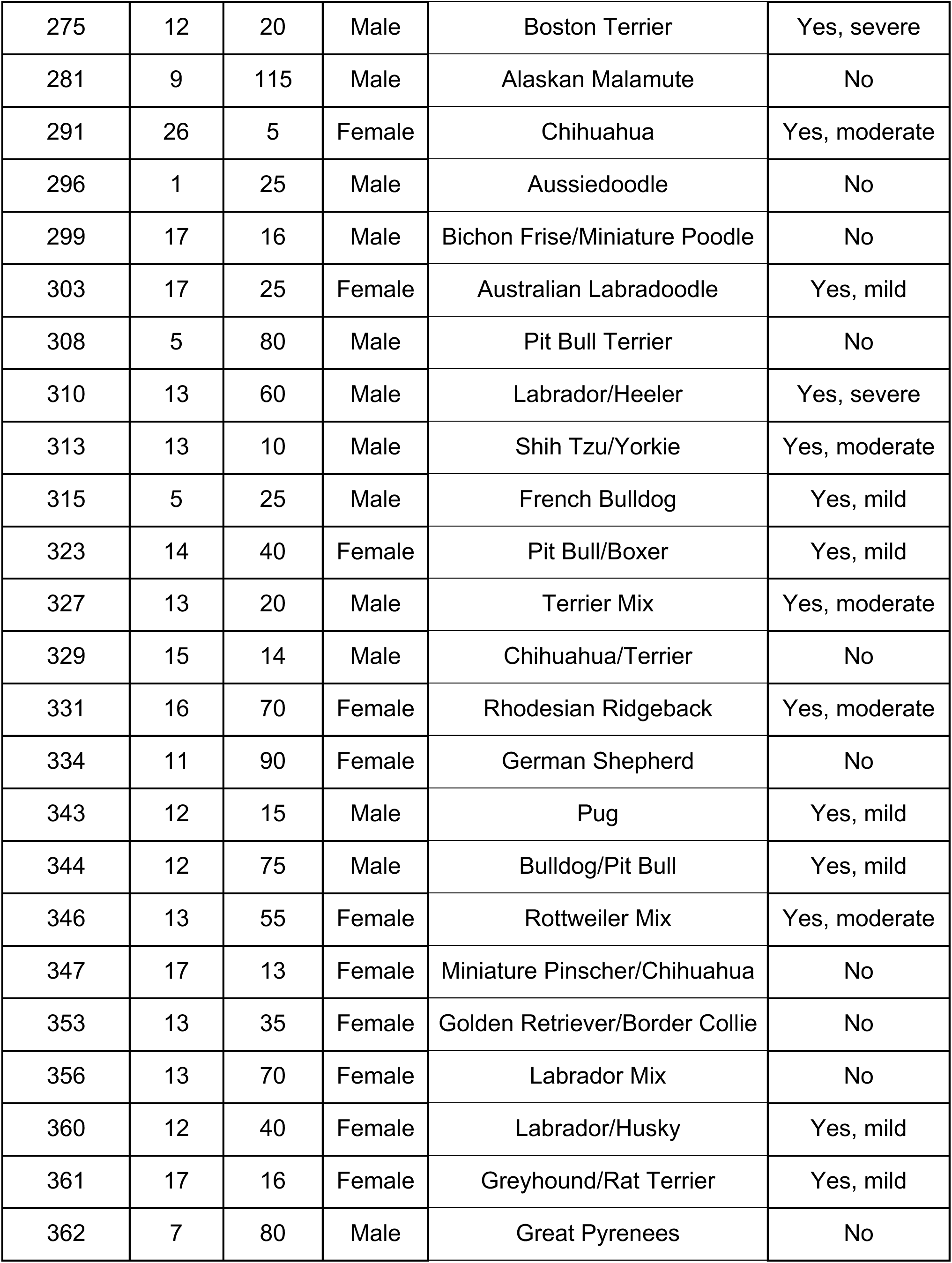
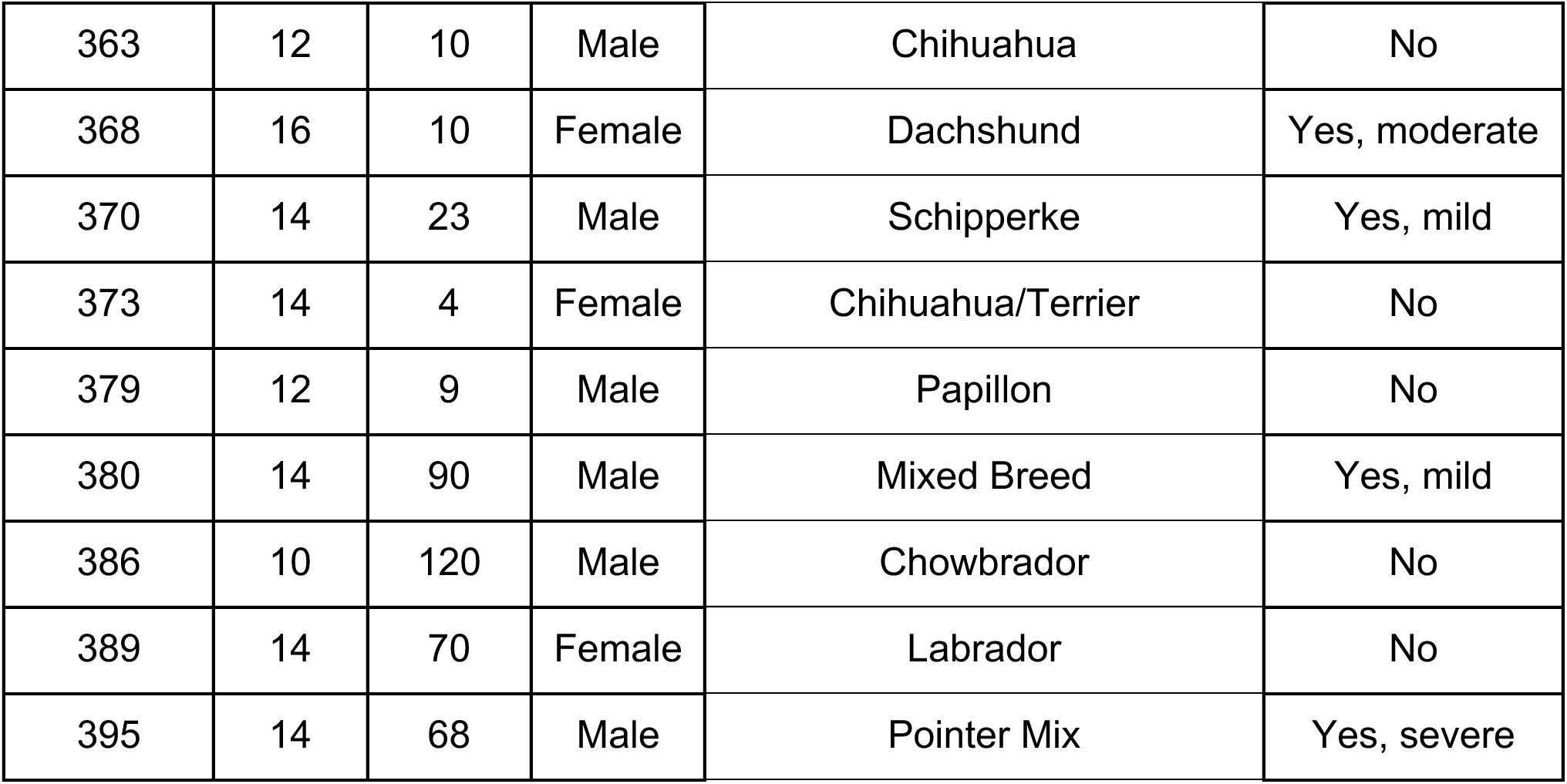
Characteristics of the canine brain bank donors. For each donor, the table lists owner-reported age, body weight, sex, breed, and signs of cognitive dysfunction in the six months prior to death. Sex is listed as unknown (Unk) where not recorded, and cognitive status is graded as no signs, mild, moderate, or severe, or listed as unknown where not reported.

Common breeds and mixes in our cohort include Chihuahuas and Chihuahua mixes, Pitbull mixes, Labrador mixes, and several small terrier breeds. Many donors are of mixed ancestry, which is indicative of the heterogeneity of the companion dog population from which our donors originated. This breed diversity is of particular interest for comparative neuroscience and aging studies, given the well-documented relationship between body size, breed, and lifespan in dogs, and the substantial variation in absolute brain size across breeds, from approximately 55 cm³ in a Pomeranian to 117 cm³ in an English Mastiff (Garamszegi et al., 2023).

Owner-reported signs of cognitive dysfunction in the six months prior to death were documented for most donors. Of the 55 donors, 27 (49.09%) had no reported signs of confusion, 27 (49.09%) had signs ranging from mild to severe, and 1 (1.82%) had unknown or unreported status.

### Perfusion quality

In n = 45 of the dogs in our brain bank, we attempted to preserve their brain via perfusion fixation (the others were fixed via immersion fixation alone). We graded perfusion quality across cases via gross examination, post-perfusion CT, and the clearance of blood vessels on histological examination (**Figure 2**). The gross examination and CT scan modalities were significantly correlated (Spearman rho = 0.66, p = 0.0009, n = 22). The histological vessel-clearance score was also positively correlated with both the gross score (rho = 0.76, p = 0.001, n = 15) and the CT score (rho = 0.70, p = 0.0017, n = 17), although these comparisons were limited by the smaller number of donors with histological grading. The consistency of these three measures corroborates our previous data using this multimodal approach to assessing perfusion quality in human brain donors (Garrood et al., 2025b).

**Figure 2.**
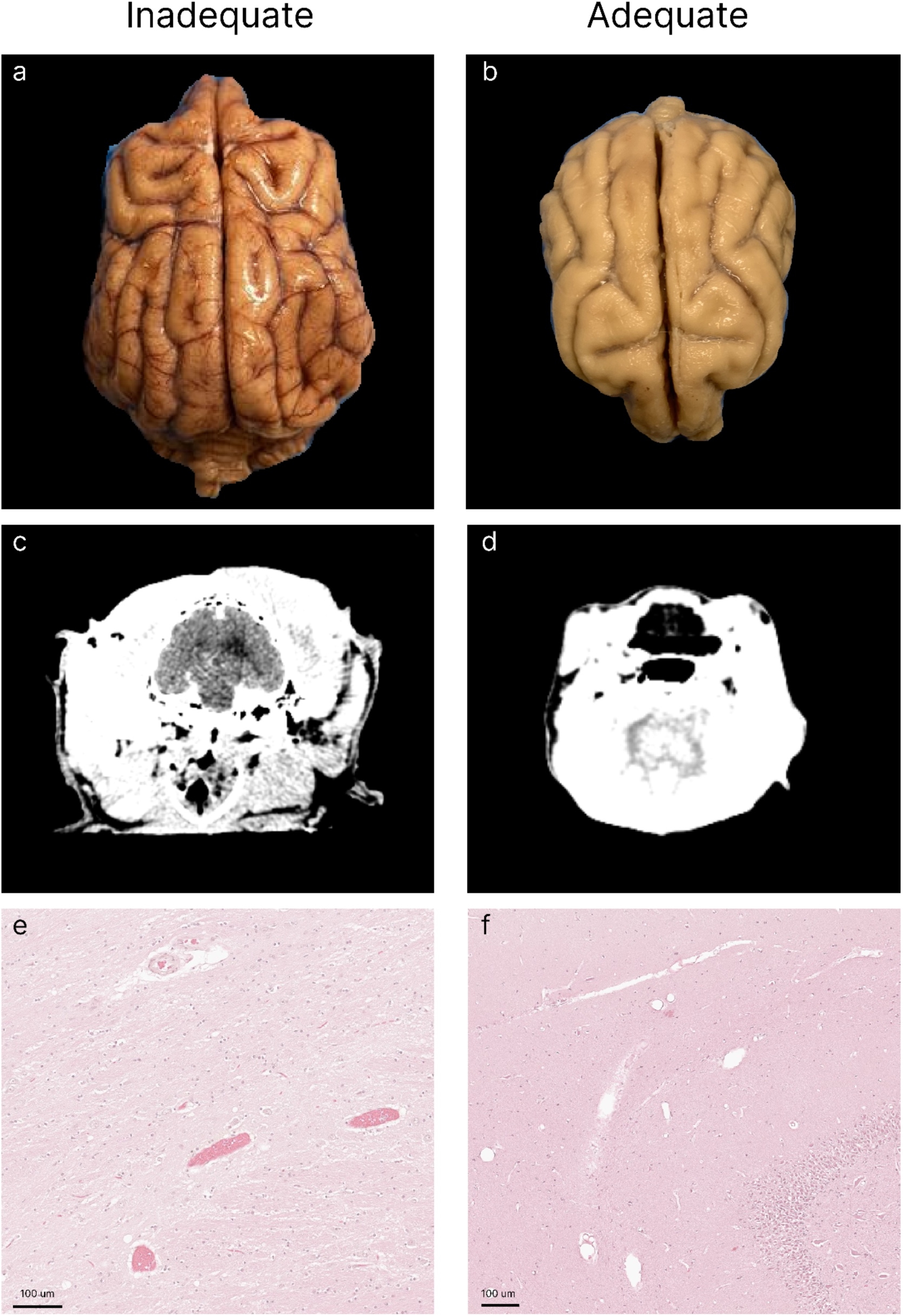
Representative examples of inadequate and adequate perfusion across imaging modalities. Each panel shows a separate case, grouped by column to illustrate the range of perfusion quality observed. The left column (**a**, **c**, **e**) shows examples graded as inadequately perfused; the right column (**b**, **d**, **f**) shows examples graded as adequately perfused. **a**, **b**: Gross examination of the dorsal brain surface immediately after brain extraction (**a**, donor 177; **b**, donor 235). Inadequate perfusion is characterized by retained blood in surface vessels and a pink to tan color, whereas adequate perfusion shows clearance of surface vessels and pale, evenly fixed tissue. **c**, **d**: Post-perfusion axial CT scans (**c**, donor 296; **d**, donor 214). **e**, **f**: Light microscopy stained with H&E (**e**, donor 296, hippocampus; **f**, donor 291, thalamus). The inadequately perfused example (**e**) shows intravascular material retained within vessels, whereas the adequately perfused example (**f**) shows clearance of intravascular material from the majority of blood vessels. Scale bars: 100 μm.

We next measured the correlations of several donor and perfusion procedure variables with perfusion quality metrics (**Figure 3**). Body weight showed the most consistent relationship, with heavier dogs tending to have lower gross perfusion scores (rho = −0.64, p = 1.3 × 10^−5^, n = 39). The average flow rate per body weight was also significantly and positively correlated with gross perfusion scores (rho = 0.40, p = 0.034, n = 29). Age, PMI, duration of perfusion, lowest and highest recorded perfusion pressure, and volume of fluid perfused were not significantly associated with the perfusion quality metrics. Notably, several of the donor characteristic variables are correlated in our cohort. For example, age and weight were found to be correlated (rho = −0.43, p = 0.0012, n = 55), with heavier dogs tending to be younger. This makes it difficult to fully separate the relationships between donor characteristics and perfusion quality in the data set that we describe here.

**Figure 3.**
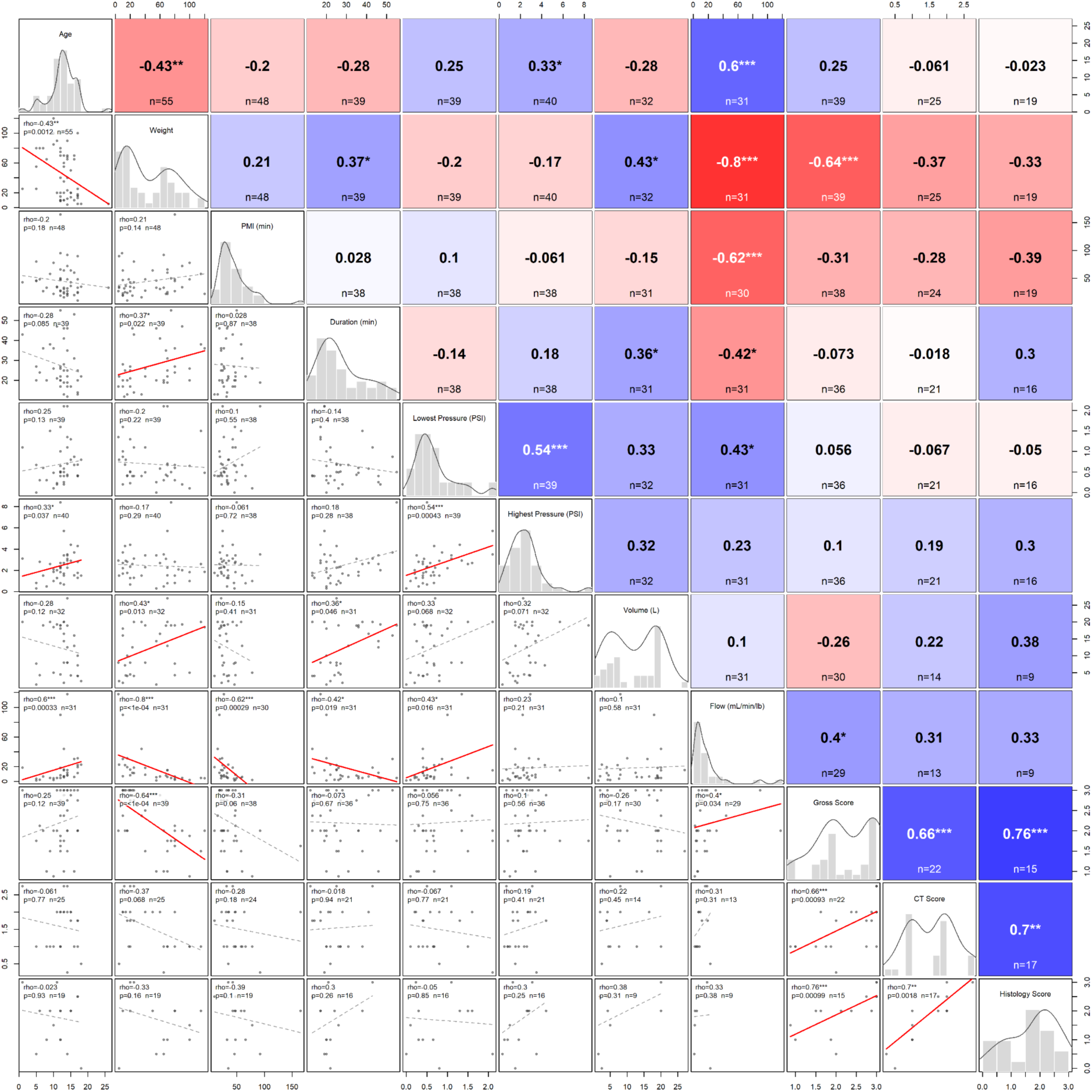
Correlations among donor characteristics, perfusion parameters, and perfusion quality metrics. Specifically, pairwise Spearman rank correlations are shown for donor age, body weight, postmortem interval (PMI), duration of perfusion, lowest recorded pressure, highest recorded pressure, perfusate volume, average flow rate per body weight (calculated as the total volume perfused per the duration of the perfusion in minutes per the weight of the animal), and the average gross, CT, and histology quality scores across brain regions in the perfused cases. The lower triangle shows scatterplots with regression lines (solid red where the correlation is significant at a nominal p-value < 0.05, dashed gray where it is not), the diagonal shows the distribution of each variable as a histogram with an overlaid kernel density estimate, and the upper triangle shows the correlation coefficient and sample size for each pair, shaded by the strength and direction of the correlation (red, negative; blue, positive). Significance is indicated as follows for nominal p-values: ***p < 0.001, **p < 0.01, *p < 0.05. Sample sizes vary by pair because not all variables were recorded for all donors.

We also note a few qualitative observations from our experience performing perfusion fixation in this context. First, we found that gravity appeared to play a substantial role in mediating where in the dog’s body the fixative solutions flowed. After we noticed this, we began tilting the table downward so that gravity would help deliver fixative to the brain, consistent with previous reports that gravity affects the regional distribution of perfusate in whole body postmortem perfusion (Nathan, 1970). Second, there was a substantial amount of anatomical variation. For example, in one case the aorta appeared to be right-sided, which has previously been reported in the context of anatomical variation in dogs (Schorn et al., 2021). Third, clots were very commonly present, presumably due to the ischemic delay before perfusion. They were uniformly of the dark red “cruor” type (Solarino et al., 2025). These clots were found to occur even within a PMI of approximately 20 minutes, broke apart easily, and were pumped out of the circulation through the cut right atrium. We did not observe the yellow “chicken fat” clots that we have seen when perfusing blood vessels in postmortem human brains (Garrood et al., 2025b). This is consistent with “chicken fat” clots arising from an agonal process or something otherwise specific to the way humans typically die, rather than from the controlled euthanasia of our donors (Solarino et al., 2025).

### Lipofuscin analysis

Lipofuscin burden was quantified in 21 donors with available histological sections from the thalamus and hippocampus (**Figure 4**). Lipofuscin density was found to have a significant rank correlation with age in both the thalamus (ρ = 0.84, p = 1.5 x 10^−6^) and the hippocampus (ρ = 0.67, p = 0.00083) (**Figure 5**). Similarly, the number of lipofuscin-positive objects had a significant rank correlation with age in both the thalamus (ρ = 0.80, p = 1.2 x 10^−5^) and the hippocampus (ρ = 0.73, p = 0.00019) (**Figure 5**). Of the 21 donors with lipofuscin data, 9 had no reported signs of cognitive dysfunction in the six months prior to death, 11 had signs ranging from mild to severe, and 1 had unknown cognitive status and was excluded from the cognitive dysfunction comparison. Dogs with cognitive dysfunction had a higher lipofuscin density in the hippocampus (Mann-Whitney U = 19, p = 0.023) and a higher hippocampal annotation count (U = 19, p = 0.023), but differences in the thalamus were not significantly different for either lipofuscin density (U = 28, p = 0.111) or annotation count (U = 27, p = 0.095) (**Figure 6**). Notably, because lipofuscin accumulation is a hallmark of brain aging, and because dogs with cognitive dysfunction in our cohort were generally older, we cannot determine from these data the extent to which the observed association between hippocampal lipofuscin and cognitive dysfunction is independent of age and other unmeasured pathology. Larger cohorts that allow controlling for chronological and biological measures of age, and other variables, would be necessary to further interrogate this question.

**Figure 4.**
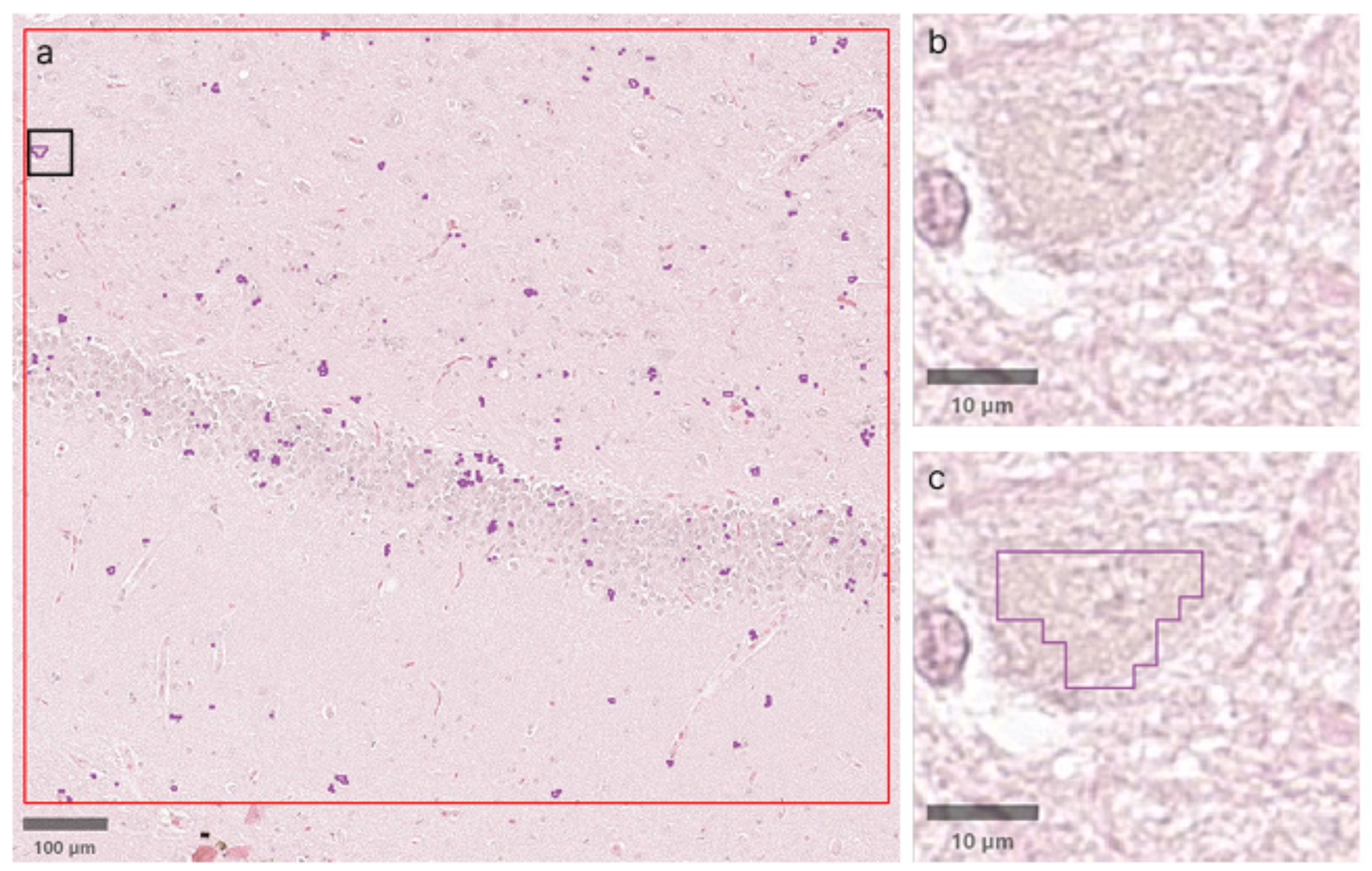
Representative lipofuscin quantification in a portion of an H&E stained whole slide image (donor 204, hippocampus sample). **a**: Overview with the analysis region outlined in red; lipofuscin-positive pixels detected by the QuPath classifier appear as scattered purple foci. The black box indicates the region magnified in (**b**) and (**c**). **b**: Higher-magnification view of neurons containing intracytoplasmic lipofuscin. **c**: The same field with the classifier-generated annotation outlined in purple, marking pixels identified as lipofuscin. Scale bars: 100 μm (**a**) and 10 μm (**b**, **c**).

**Figure 5.**
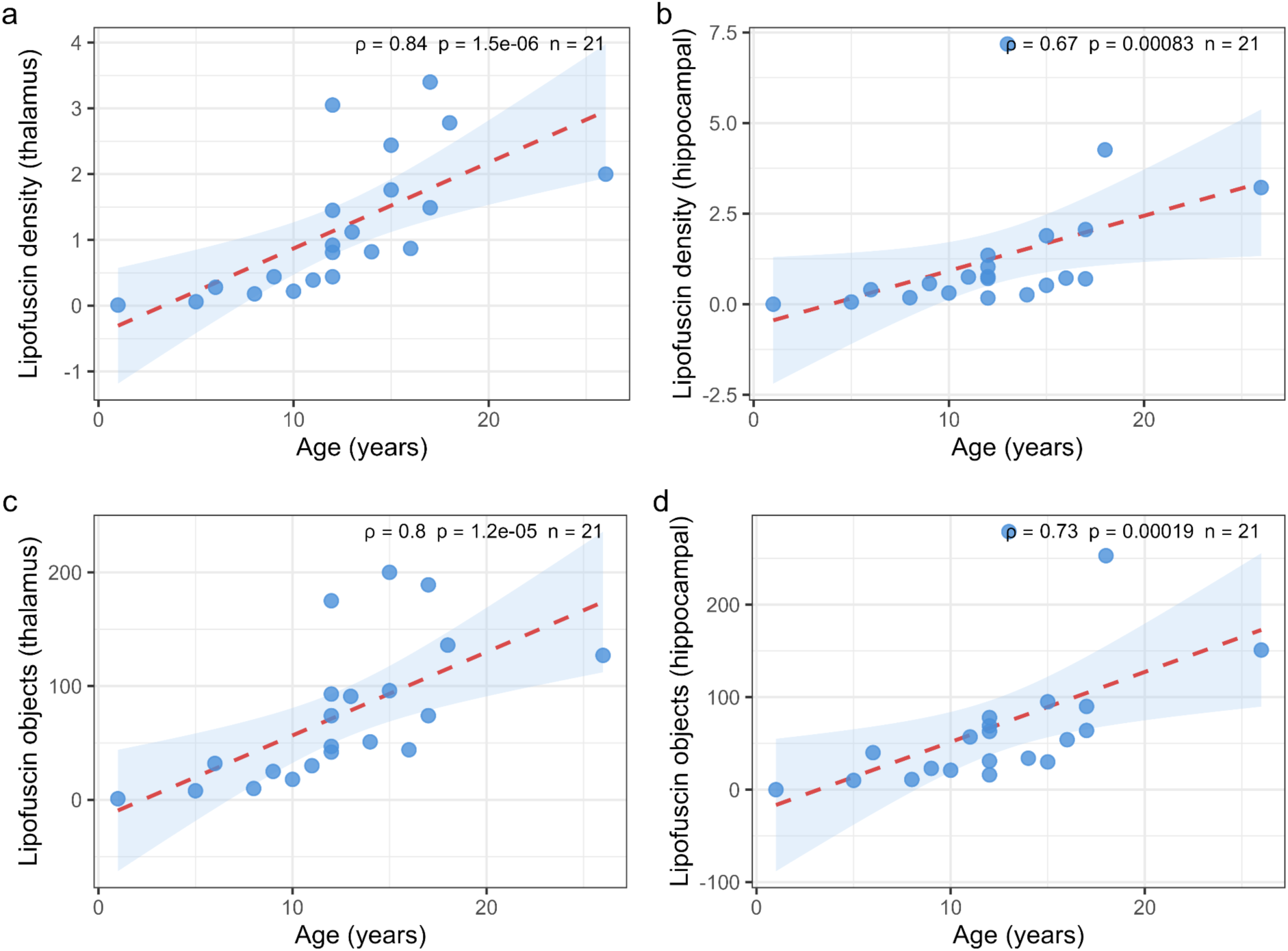
Lipofuscin accumulation increases with age in the canine brain. Lipofuscin density (**a**, **b**) and lipofuscin-positive object count (**c**, **d**) are plotted against age for the thalamus (**a**, **c**) and hippocampus (**b**, **d**). Red dashed lines indicate linear regression lines for visualization only with 95% confidence intervals (blue shading). Spearman rank correlations are shown for each panel (n = 21 donors).

**Figure 6.**
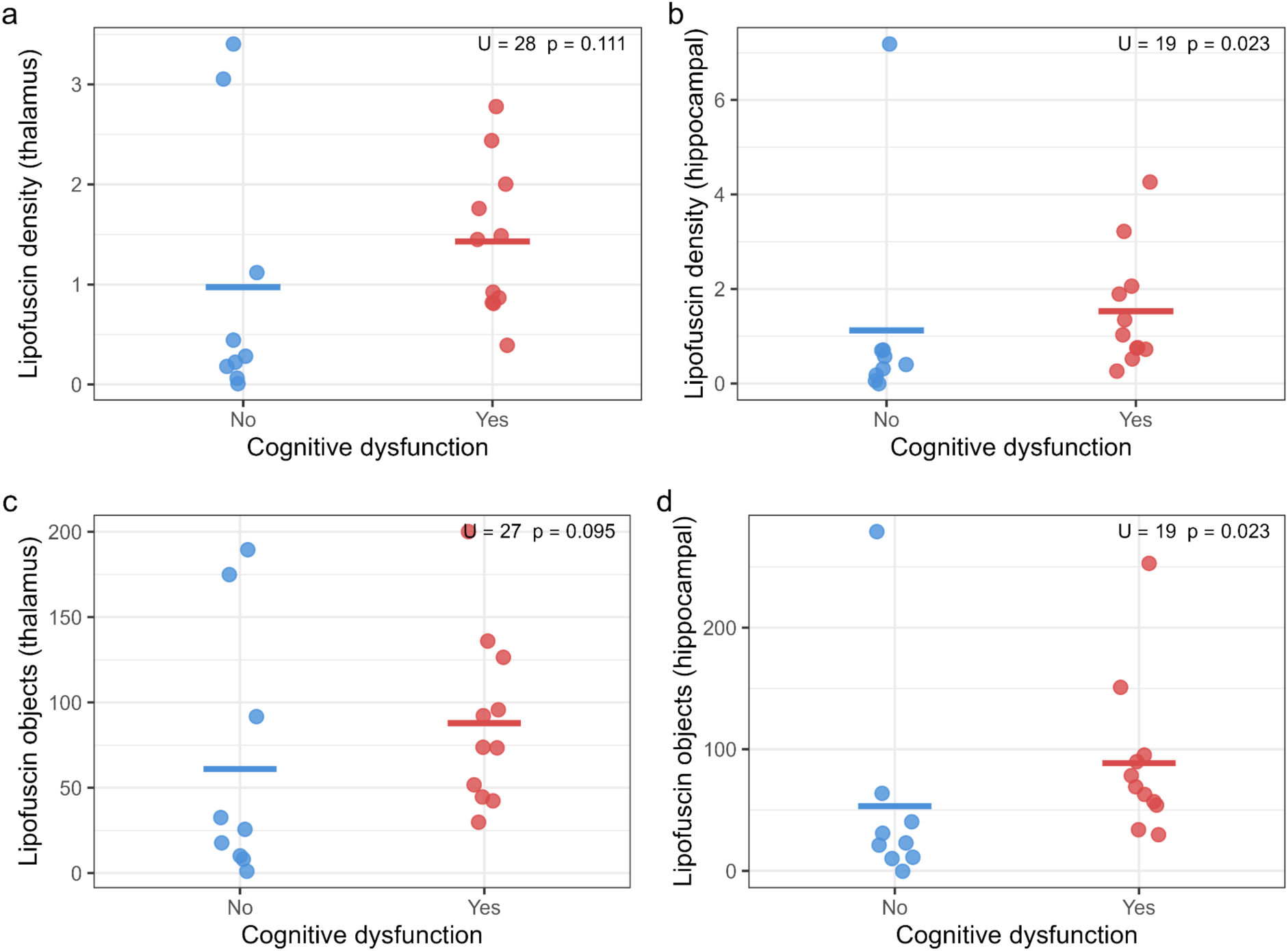
Hippocampal lipofuscin burden is elevated in dogs with owner-reported cognitive dysfunction. Lipofuscin density (**a**, **b**) and lipofuscin-positive object count (**c**, **d**) are compared between dogs with no reported cognitive dysfunction (blue, n = 9) and dogs with owner-reported signs of cognitive dysfunction in the six months prior to death (red, n = 11). Horizontal bars indicate group means. Mann-Whitney U test statistics and p-values are shown for each panel.

### Preliminary ultrastructural assessment

To evaluate whether perfusion fixed tissue from our canine brain bank is suitable for ultrastructural analysis, we performed transmission electron microscopy on cortical samples from two donors (**Figure 7**). Major cellular structures were readily identifiable in images from both donors, including nuclei, mitochondria with discernible cristae, myelinated axons, and synapses. Myelin lamellae were frequently disrupted, which is a well-known artifact of conventional aldehyde fixation and EM processing, even in non-ischemic tissue (Anderson et al., 2014; Möbius et al., 2016). Some degree of membrane-bound unstained areas were observed across nearly all images examined. Such unstained areas of this kind are generally attributed to ischemic fluid shifts and associated swelling of cellular processes prior to fixation, especially of astrocytes (Kalimo et al., 1977; Krassner et al., 2023; Garrood et al., 2025a). At higher magnification, synaptic profiles with visible postsynaptic densities and presynaptic vesicle pools could also be identified. These preliminary observations, based on a limited number of images from two donors, suggest that at least a subset of the perfusion fixed tissue from this canine brain bank is amenable to ultrastructural investigation. The conditions under which the tissue is sufficiently preserved for automated connectome tracing will require substantially more data and evaluation.

**Figure 7.**
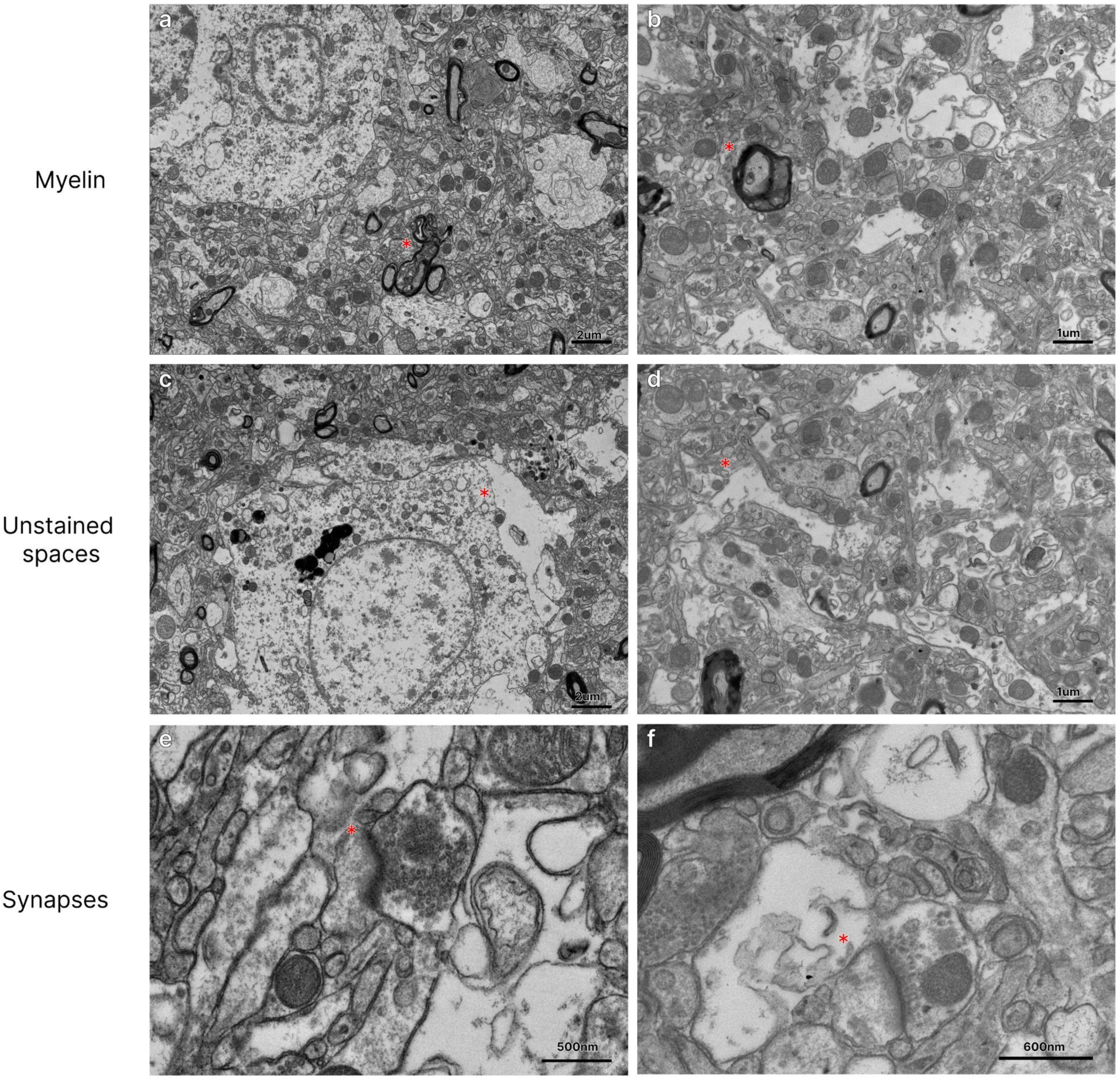
Representative electron microscopy images from two perfusion fixed donors, dog 65 (**a**, **c**, **e**) and dog 177 (**b**, **d**, **f**). Rows show myelinated axons (**a**, **b**), unstained spaces (**c**, **d**), and synapses (**e**, **f**). Red stars are placed just to the left of structures of interest in each panel: a myelinated axon (**a**, **b**), a membrane-bound unstained area (**c**, **d**), and a synapse (**e**, **f**). Scale bars: 2 μm (**a**, **c**), 1 μm (**b**, **d**), 500 nm (**e**), 600 nm (**f**).

## Discussion

In this manuscript, we have described the development of our perfusion fixation-based canine brain bank. We found that aortic cannulation yields high-quality perfusion in some but not all cases. In our preliminary histologic investigations, we found that hippocampal lipofuscin burden is elevated in dogs with cognitive dysfunction. To the best of our knowledge, this is the first quantitative assessment of the association between lipofuscin burden on histology and cognitive dysfunction status in the domestic dog. However, we note that this histological analysis of lipofuscin is preliminary and exploratory. The light and electron microscopy data we described in this manuscript is already publicly available (see Data Availability). We plan to make tissue from this brain bank available to the neuroscience research community upon request from qualified investigators. Below, we describe future considerations and plans for our canine brain bank.

One of the most notable features of the domestic dog as a model for brain aging research is the dramatic variation in the typical lifespan across breeds. This ranges from approximately 7 years in giant breeds such as the Irish Wolfhound up to 16 years in small breeds such as the Papillon, with weight being the strongest predictor of lifespan across breeds (Greer et al., 2007). This raises the question of whether breed-specific lipofuscin accumulation rate is also associated with breed-specific lifespan, and whether lipofuscin density could serve as a readout of biological aging to capture differences in the rate of brain cell aging between breeds. More broadly, it is unclear the extent to which lipofuscin is a passive correlate of aging or actively contributes to neuronal dysfunction, for example by impairing lysosomal function and proteostasis (Brunk and Terman, 2002). As our canine brain bank grows, the tissue could be used to help adjudicate between these possibilities, for example by measuring whether lipofuscin density covaries with other markers of age-related neuronal or lysosomal dysfunction.

The variable perfusion quality we observed is indicative of the challenges of achieving rapid whole brain fixation in dogs. Several factors likely contribute to this, including differences in flow rate to body size ratio, vascular anatomy, and the PMI prior to perfusion. Improving perfusion consistency is a priority, because the quality of ultrastructural preservation determines what downstream analyses are possible. If fixation quality can be made reliably high, canine brain tissue could become amenable to connectomic analysis (Lichtman et al., 2014; Mikula, 2016; Helmstaedter, 2026). Dogs are an appealing model for comparative connectomics, because they possess complex social cognition, yet have brains far smaller than human brains, making large-scale reconstruction more theoretically feasible. One advantage of our approach is that the controlled euthanasia setting provides short PMIs compared to most human brain banks, which could be leveraged for systematic studies of how PMI, age, vascular disease, and other naturalistic variables affect perfusion quality and ultrastructural preservation. What combination of PMI, perfusion pressure, perfusate composition, and other parameters is needed to consistently achieve connectomics-grade preservation in a canine donor program such as ours is an open question.

This study has several limitations. First, the preliminary cohort that we describe here is relatively small. Expanding the cohort is an obvious priority, as the research value of a brain bank depends in large part on the size and diversity of its collection (Wang et al., 2019). Second, cognitive dysfunction status was based on a single question about recent signs of confusion, which is a coarse measure of a multidimensional syndrome. In future work, we plan to use validated owner assessment instruments such as the Canine Cognitive Dysfunction Rating scale questionnaire to improve phenotyping (Salvin et al., 2011). Third, we did not perform immunostaining in the present study, such as for well known markers of cognitive dysfunction, such as amyloid beta or tau. Finally, the perfusion protocols evolved over the course of the study as we optimized our methods, which helped us to identify the most effective methods, but introduced a significant confounding variable when comparing perfusion quality across cases.

## Conclusions

We have described the methods for our canine brain bank, in which we primarily use perfusion fixation for initial preservation. We have also presented preliminary analyses of brain aging and ultrastructure that are intended to demonstrate its future research value. Looking ahead, we expect that the value of this resource will grow with the size, diversity, and preservation quality of the collection. For example, a larger cohort would allow the relationship between hippocampal lipofuscin and cognitive dysfunction to be disentangled from chronological age. More generally, it would allow us to ask whether breeds with longer lifespans also have slower brain aging, as measured by markers such as lipofuscin. We hope this resource and the methods we present will help support more research on the domestic dog, donated in an ethical manner after death for unrelated causes, in the fields of neuroscience and aging.

## Abbreviations

ACA: Anterior cerebral artery
CT: Computed tomography
EM: Electron microscopy
H&E: Hematoxylin and eosin
LM: Light microscopy
MCA: Middle cerebral artery
NBF: Neutral buffered formalin
PCA: Posterior cerebral artery
PEG35: Polyethylene glycol 35 kDa
PMI: Postmortem interval
WSI: Whole slide image.

## Author Contributions

A.T.M., K.F., and J.F.C. conceptualized the study. S.D., A.B., M.G., A.S., A.P., and L.P. performed laboratory experiments and data analysis. A.T.M. wrote the initial draft of the manuscript. All authors reviewed the manuscript and approved the final manuscript.

## Data Availability

The light and electron microscopy data described in this manuscript are publicly available at the following DOIs: https://doi.org/10.5281/zenodo.21047579, https://doi.org/10.5281/zenodo.15312788, and https://doi.org/10.5281/zenodo.18805394. The QuPath pixel classifier used for lipofuscin quantification is available at this DOI: https://doi.org/10.5281/zenodo.21225320. Code and data to reproduce the plots and analysis is available at https://github.com/andymckenzie/Canine_brain_bank.

## Acknowledgments

This work is dedicated to the memory of Dr. Amy Pelton, a partner veterinarian and valued collaborator.

## Declaration of Generative AI Technologies

In the preparation of this manuscript, the authors used Claude (Anthropic) for both programming assistance and to improve the manuscript’s language. All AI tool-assisted content was reviewed and edited by the authors, who take full responsibility for the final publication.

## References

Anderson, A.J., Hooshmand, M.J., Cummings, B.J., 2014. Improved pre-embedded immuno-electron microscopy procedures to preserve myelin integrity in mammalian central nervous tissue, in: Microscopy: Advances in Scientific Research and Education. Formatex Research Center, pp. 59–65.

Barton, S.A., Smaers, J.B., Serpell, J.A., Hecht, E.E., 2025. Brain-Behavior Differences in Premodern and Modern Lineages of Domestic Dogs. J Neurosci 45, e2032242025. 10.1523/JNEUROSCI.2032-24.2025

Bodian, D., 1936. A new method for staining nerve fibers and nerve endings in mounted paraffin sections. The Anatomical Record 65, 89–97. 10.1002/ar.1090650110

Brunk, U.T., Terman, A., 2002. Lipofuscin: mechanisms of age-related accumulation and influence on cell function. Free Radic Biol Med 33, 611–619. 10.1016/s0891-5849(02)00959-0

Camstra, K.M., Srinivasan, V.M., Collins, D., Chen, S., Kan, P., Johnson, J., 2020. Canine Model for Selective and Superselective Cerebral Intra-Arterial Therapy Testing. Neurointervention 15, 107–116. 10.5469/neuroint.2020.00150

Cardy, T.J.A., Jewth-Ahuja, D., Crawford, A.H., 2022. Perceptions and attitudes towards companion animal brain banking in pet owners: A UK pilot study. Vet Rec Open 9, e36. 10.1002/vro2.36

Cummings, B.J., Head, E., Ruehl, W., Milgram, N.W., Cotman, C.W., 1996. The canine as an animal model of human aging and dementia. Neurobiol Aging 17, 259–268. 10.1016/0197-4580(95)02060-8

Eden, A.R., Correia, M.J., 1981. Improved fixation of the pigeon brain by transcardiac carotid catheterization. Physiol Behav 27, 947–949. 10.1016/0031-9384(81)90066-4

Few, A., Getty, R., 1967. Occurrence of lipofuscin as related to aging in the canine and porcine nervous system. J Gerontol 22, 357–368. 10.1093/geronj/22.3.357

Gage, G.J., Kipke, D.R., Shain, W., 2012. Whole animal perfusion fixation for rodents. J Vis Exp. 10.3791/3564

Garamszegi, L.Z., Kubinyi, E., Czeibert, K., Nagy, G., Csörgő, T., Kolm, N., 2023. Evolution of relative brain size in dogs-no effects of selection for breed function, litter size, or longevity. Evolution 77, 1591–1606. 10.1093/evolut/qpad063

Garrood, M., Keberle, A., Sowa, A., Janssen, W., Thorn, E.L., Sanctis, C.D., Farrell, K., Crary, J.F., McKenzie, A.T., 2025a. Evaluating ultrastructural preservation quality in banked brain tissue. Free Neuropathol 6, 13. 10.17879/freeneuropathology-2025-6763

Garrood, M., Keberle, A., Taylor, G.A., Thorn, E.L., Sanctis, C.D., Farrell, K., Crary, J.F., Sparks, J.S., McKenzie, A.T., 2025b. Mechanical perfusion in brain banking: methods of assessment and relationship to the postmortem interval. Free Neuropathol 6, 20. 10.17879/freeneuropathology-2025-8880

Greer, K.A., Canterberry, S.C., Murphy, K.E., 2007. Statistical analysis regarding the effects of height and weight on life span of the domestic dog. Res Vet Sci 82, 208–214. 10.1016/j.rvsc.2006.06.005

Head, E., 2013. A canine model of human aging and Alzheimer’s disease. Biochim Biophys Acta 1832, 1384–1389. 10.1016/j.bbadis.2013.03.016

Helmstaedter, M., 2026. Synaptic-resolution connectomics: towards large brains and connectomic screening. Nat Rev Neurosci 27, 101–120. 10.1038/s41583-025-00998-z

Jewell, P.A., 1952. The anastomoses between internal and external carotid circulations in the dog. J Anat 86, 83–94.

Kalimo, H., Garcia, J.H., Kamijyo, Y., Tanaka, J., Trump, B.F., 1977. The ultrastructure of “brain death”. II. Electron microscopy of feline cortex after complete ischemia. Virchows Arch B Cell Pathol 25, 207–220.

Krassner, M.M., Kauffman, J., Sowa, A., Cialowicz, K., Walsh, S., Farrell, K., Crary, J.F., McKenzie, A.T., 2023. Postmortem changes in brain cell structure: a review. Free Neuropathol 4, 4–10. 10.17879/freeneuropathology-2023-4790

Lee, M.C., Reid, I.A., Ramsay, D.J., 1986. Blood flows in the maxillocarotid anastomoses and internal carotid artery of conscious dogs. Anat Rec 215, 192–197. 10.1002/ar.1092150212

Lichtman, J.W., Pfister, H., Shavit, N., 2014. The big data challenges of connectomics. Nat Neurosci 17, 1448–1454. 10.1038/nn.3837

McEnhill, R., Borghese, H., Moore, S.A., 2024. Pet owner perspectives, motivators and concerns about veterinary biobanking. Front Vet Sci 11, 1359546. 10.3389/fvets.2024.1359546

McFadden, W.C., Walsh, H., Richter, F., Soudant, C., Bryce, C.H., Hof, P.R., Fowkes, M., Crary, J.F., McKenzie, A.T., 2019. Perfusion fixation in brain banking: a systematic review. Acta Neuropathol Commun 7, 146. 10.1186/s40478-019-0799-y

McGrath, S., Hull, E., Dunbar, M.D., Prescott, J., Keyser, A.J., MacLean, E., Darvas, M., Latimer, C., Moreno, J., MacCoss, M.J., Kauffman, M., Litwin, P., Castelhano, M., Kaeberlein, M., Keene, C.D., Dog Aging Project Consortium, 2025. The companion dog as a translational model for Alzheimer’s disease: Development of a longitudinal research platform and post mortem protocols. Alzheimers Dement 21, e70630. 10.1002/alz.70630

McKenzie, A.T., Nnadi, O., Slagell, K.D., Thorn, E.L., Farrell, K., Crary, J.F., 2024. Fluid preservation in brain banking: a review. Free Neuropathol 5, 5–10. 10.17879/freeneuropathology-2024-5373

Mikula, S., 2016. Progress Towards Mammalian Whole-Brain Cellular Connectomics. Frontiers in Neuroanatomy 10.

Möbius, W., Nave, K.-A., Werner, H.B., 2016. Electron microscopy of myelin: Structure preservation by high-pressure freezing. Brain Res 1641, 92–100. 10.1016/j.brainres.2016.02.027

Nanda, B.S., Getty, R., 1973. Occurrence of aging pigment (lipofuscin) in the nuclei and cortices of the canine brain. Experimental Gerontology 8, 1–7. 10.1016/0531-5565(73)90044-2

Nathan, H., 1970. A simple method of embalming human cadavers by intracardiac injection. Acta Anat (Basel) 77, 155–159. 10.1159/000143538

Nesic, S., Vučićević, I., Marinković, D., Kukolj, V., Aničić, M., Ristoski, T., Nikolić, S., Aleksić-Kovačević, S., 2021. Histochemical characteristics and distribution of lipofuscin and polyglucosan bodies in the brain of dogs more than 10 years old. Veterinarski Glasnik 75, 57–68. 10.2298/VETGL201002001N

Park, S.-G., Kang, M.-H., Lee, C.-M., Sur, J., Park, H.-M., 2016. Cognitive Dysfunction Syndrome with Lipofuscinosis in a Maltese Dog. Pakistan Veterinary Journal 36, 508–510.

Salt, C., Morris, P.J., German, A.J., Wilson, D., Lund, E.M., Cole, T.J., Butterwick, R.F., 2017. Growth standard charts for monitoring bodyweight in dogs of different sizes. PLoS One 12, e0182064. 10.1371/journal.pone.0182064

Salvin, H.E., McGreevy, P.D., Sachdev, P.S., Valenzuela, M.J., 2011. The canine cognitive dysfunction rating scale (CCDR): a data-driven and ecologically relevant assessment tool. Vet J 188, 331–336. 10.1016/j.tvjl.2010.05.014

Sándor, S., Czeibert, K., Salamon, A., Kubinyi, E., 2021. Man’s best friend in life and death: scientific perspectives and challenges of dog brain banking. Geroscience 43, 1653–1668. 10.1007/s11357-021-00373-7

Schorn, C., Hildebrandt, N., Schneider, M., Schaub, S., 2021. Anomalies of the aortic arch in dogs: evaluation with the use of multidetector computed tomography angiography and proposal of an extended classification scheme. BMC Vet Res 17, 387. 10.1186/s12917-021-03101-7

Solarino, B., Ambrosi, L., Benevento, M., Ferorelli, D., Buschmann, C., Nicolì, S., 2025. Cadaver clots: a systematic review of the literature. Forensic Sci Med Pathol 21, 1831–1842. 10.1007/s12024-025-00976-y

Trahanas, J., Hoffman, J., McMaster, W., Tipograf, Y., Absi, T., Balsara, K., Shah, A., 2022. DCD Organ Procurement with Normothermic Regional Perfusion. CTSNet. 10.25373/ctsnet.19699936

Wahyudi, D.P., Makkiyah, F.A., Buana, G.R., 2025. A comparative analysis of transcardial perfusion techniques for brain fixation in rodents: A call for standardization of transcardial perfusion. The Health Researcher’s Journal 2, 7–13. 10.00000/6nrjth51

Wang, L., Xia, Y., Chen, Y., Dai, R., Qiu, W., Meng, Q., Kuney, L., Chen, C., 2019. Brain Banks Spur New Frontiers in Neuropsychiatric Research and Strategies for Analysis and Validation. Genomics Proteomics Bioinformatics 17, 402–414. 10.1016/j.gpb.2019.02.002

Whiteford, R., Getty, R., 1966. Distribution of lipofuscin in the canine and porcine brain as related to aging. J Gerontol 21, 31–44. 10.1093/geronj/21.1.31

Wolf, M., Beck, A., Paredes, L., Darcy, S., Parra, A., Taylor, G.A., Garrood, M., Thorn, E.L., Sanctis, C.D., Crary, J.F., Farrell, K., McKenzie, A.T., 2026. Brain extraction for fixed tissue banking: a technical report. Free Neuropathol 7, 11. 10.17879/freeneuropathology-2026-9411

